# A subpopulation of Talin 1 resides in the nucleus and regulates gene expression

**DOI:** 10.1101/2022.03.15.484419

**Authors:** Alejandro J. Da Silva, Hendrik S. E. Hästbacka, Mikael C. Puustinen, Jenny C. Pessa, Benjamin T. Goult, Guillaume Jacquemet, Eva Henriksson, Lea Sistonen

**Affiliations:** Faculty of Science and Engineering, Cell Biology, Åbo Akademi University, 20520 Turku, Finland; Turku Bioscience Centre, University of Turku and Åbo Akademi University, 20520 Turku, Finland; Turku Bioimaging University of Turku and Åbo Akademi University, 20520 Turku, Finland; School of Biosciences, University of Kent, Canterbury CT2 7NJ, Kent, UK

**Keywords:** Chromatin, gene expression, nucleus, talin 1, TLN1

## Abstract

Talin 1 (TLN1) is best known for its role at focal adhesions, where it activates β-integrin receptors and transmits mechanical stimuli to the actin cytoskeleton. Interestingly, the localization of TLN1 is not restricted to the focal adhesions, but its function in other cellular compartments remains poorly understood. By utilizing both biochemical and confocal microscopy analyses, we show that TLN1 localizes to the nucleus and that it strongly interacts with the chromatin. Importantly, depletion of endogenous TLN1 results in extensive changes in the gene expression profile of human breast epithelial cells. To determine the impact of nuclear TLN1 on gene regulation, we expressed a TLN1 fusion protein containing a nuclear localization signal. Our results revealed that nuclear TLN1 regulates a specific subset of the TLN1-dependent genes. Taken together, we show that apart from localizing at the plasma membrane and cytoplasm, TLN1 also resides in the nucleus where it functions in the regulation of gene expression.

## Introduction

Accumulating evidence indicates active communication between the cell cortex and the nucleus by nuclear translocation of proteins (Zheng & Jiang, 2022). Many of these proteins are associated with transmembrane adhesion receptors, which connect the extracellular matrix (ECM) and surface proteins of neighboring cells to the cytoskeleton (Hintermann & Christen, 2019; Zheng & Jiang, 2022). These cell-ECM and cell-cell junctions provide cells with structural and mechanical stability, and both types of junctions act as signaling platforms that collect information from the extracellular space. Therefore, identifying proteins that mediate the communication between the cell cortex and the nucleus is fundamental to understand how cells respond to their surroundings. In a recent preprint article Byron and collaborators performed proteomic analyses and found that a considerable number of adhesion complex-associated proteins also reside in the nucleus (preprint: Byron *et al*, 2021). However, the functions of most adhesion-associated proteins inside the nucleus remain poorly understood.

Talin 1 (TLN1) is a 270 kDa adaptor protein that is best known for its role in focal adhesion (FA) assembly (Gough & Goult, 2018). FAs are structures characterized by proteins that crosslink actin filaments to the integrin transmembrane receptors (Case & Waterman, 2015). To achieve its function at the FAs, TLN1 has a particular domain structure that is composed of an N-terminal FERM (4.1 protein, ezrin, radixin, moesin) domain, known as the head domain, which is coupled to a flexible rod domain comprised of 13 helical bundles (Goult *et al*, 2013b). The head domain interacts with the cytoplasmic tails of β-integrin subunits, whereas the rod domain binds to actin filaments and acts as a mechanosensitive signaling hub (Goult *et al*, 2021) (Fig 1A, left panel). Dysregulation of TLN1 is associated with different diseases, such as cancer, cardiovascular malfunction, and hematologic disorders, which makes TLN1 a relevant protein in the context of therapeutics and diagnostics (Azizi *et al*, 2021; Li *et al*, 2021; Haining *et al*, 2016). Interestingly, TLN1 is not confined to the FA complexes indicating that this protein has other roles apart from the FA assembly. For example, TLN1 is localized at invadopodia, which are actin-rich protrusions that mediate cancer cell invasion and metastasis, where it acts as a scaffold to recruit the sodium/hydrogen exchanger 1 protein (Beaty *et al*, 2014). Moreover, TLN1 can also reside in the cytoplasm where it adopts a conformation that is unable to mediate the connection between integrins and the actin cytoskeleton, and therefore might have a role in other signaling pathways (Haage *et al*, 2018; Goult *et al*, 2013a). Consequently, characterizing the subcellular localization and function of TLN1 is fundamental for understanding its role in physiological processes and pathological states.

**Figure 1:**
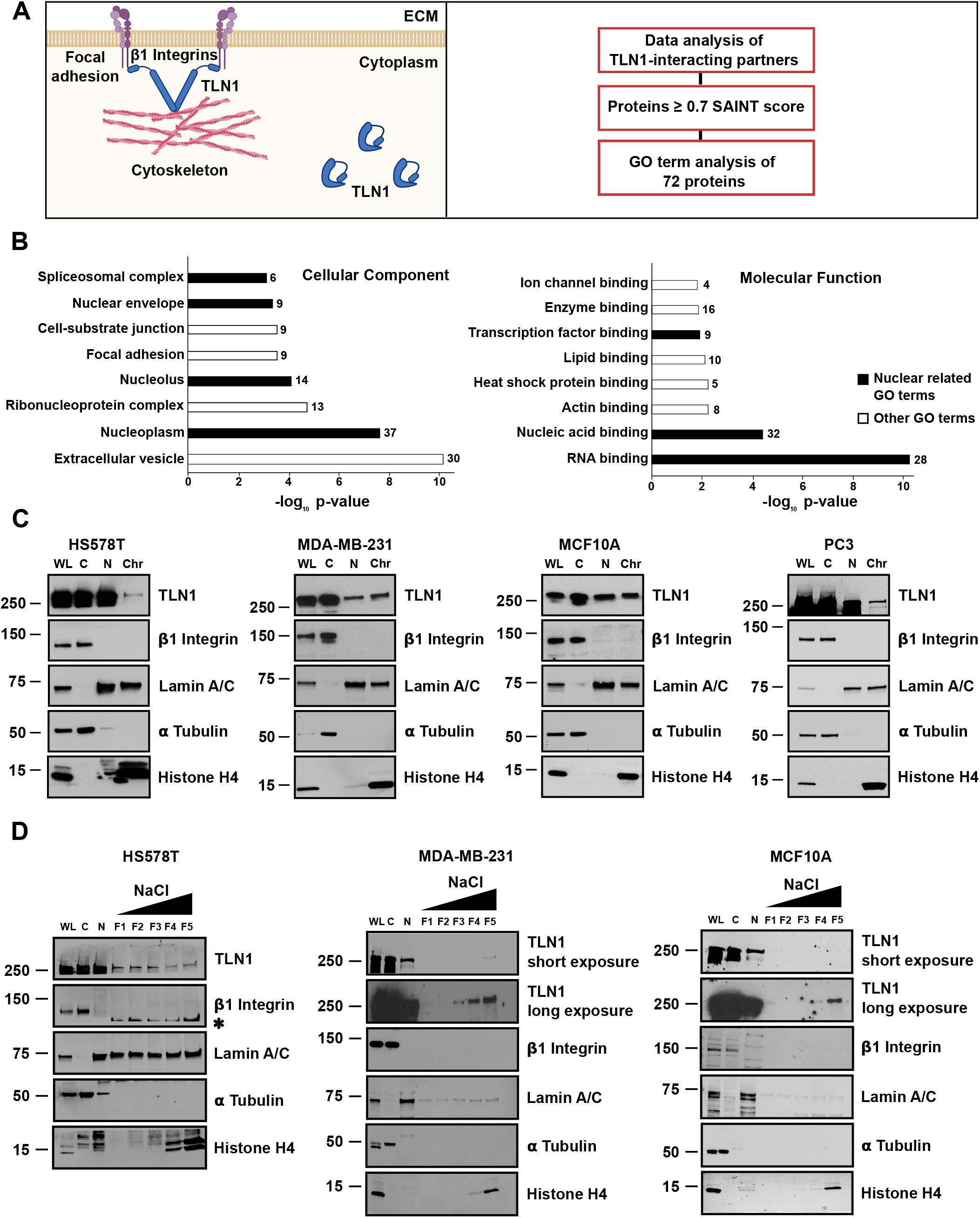
TLN1 is localized in the nucleus where it strongly interacts with the chromatin. **A)** A schematic overview of the subcellular localization of Talin 1 (TLN1) according to the current literature. TLN1 is localized in the focal adhesions and the cytoplasm (left panel). Analysis of TLN1 interacting partners previously identified by Gough and collaborators (right panel) (Gough *et al*, 2021). Made in BioRender.com **B)** Gene ontology (GO) terms associated with TLN1 interacting partners were analyzed with the online application ShinyGO (Ge *et al*, 2020). The GO terms within the “cellular component” and “molecular function” ontologies were ranked according to their p-values and the redundant terms were withdrawn. The number of proteins associated with each term is indicated, and terms composed of less than four proteins are not shown. GO terms related to the nucleus are highlighted in black. **C)** Immunoblot analysis of TLN1 in subcellular fractionations of HS578T, MDA-MB-231, MCF10A, and PC3 cells. WL: whole cell lysate, C: cytoplasmic fraction, N: nuclear fraction, Chr: chromatin fraction. To monitor the purity of the fractionation protocol the following controls were used: β1 Integrin (plasma membrane), Lamin A/C (nucleus), α Tubulin (cytoplasm), Histone H4 (chromatin). **D)** Immunoblot analysis of TLN1 in differential salt fractionation of HS578T, MDA-MB-231, and MCF10A cells. F1: 0.3, F2: 0.45, F3: 0.6, F4: 0.8, and F5: 1.2 M of NaCl. The fractionation controls were the same as in panel C and remanent signal from a previous LAM A/C immunoblot is indicated with an asterisk (*).

In this study, we investigated the subcellular localization of TLN1. Biochemical and confocal microscopy analyses showed, to our surprise, that TLN1 can also localize to the nuclei of human epithelial cells, where it strongly interacts with the chromatin. Moreover, depletion of TLN1 resulted in extensive changes in the gene expression profile of breast epithelial cells, causing upregulation and downregulation of approximately 300 and 400 genes, respectively. To determine the importance of nuclear TLN1 in gene regulation, we generated a fusion protein composed of the full-length human TLN1 coupled to GFP and a nuclear localization signal (NLS). By ectopically expressing the TLN1-NLS fusion protein, we demonstrate that enriching TLN1 in the nucleus impacts the expression of a specific subset of genes. Taken together, this study identifies TLN1 as a nuclear protein that regulates gene expression.

## Results

### TLN1 is localized in the nucleus where it strongly interacts with the chromatin

To initiate the study on the subcellular localization of talin 1 (TLN1), we examined the gene ontology (GO) terms associated with 1304 TLN1-interacting proteins, that were identified in a recently published mass spectrometry screen (Gough *et al*, 2021) (Fig 1A, right panel). Gough and collaborators determined the probability of *bona fide* protein-protein interactions by calculating the Significance Analysis of INTeractome (SAINT) score, and a total of 72 TLN1-interacting partners, which had a SAINT score ≥0.7, were chosen for our GO term analysis. Interestingly, the analysis revealed that a majority, i.e. 37, of these 72 proteins are nuclear, and 32 proteins are involved in nucleic acid binding (Fig 1B) (Table1). This unexpected result prompted us to investigate whether TLN1 is localized in the nucleus of transformed (HS578T, MDA-MB-231) and non-transformed (MCF10A) human breast epithelial cells, as well as transformed human prostate epithelial cells (PC3). To achieve fractions of high purity, we utilized a protocol based on stepwise cell lysis where the cytoplasm is separated from the intact nucleus, which in turn is lysed separately to extract proteins located either in the nucleoplasm or in the chromatin. To assess the purity of the subcellular fractions, we monitored the localization of proteins that are known to reside in the following compartments: plasma membrane (β1 integrin), cytoplasm (α tubulin), nucleus (lamin A/C), and chromatin (histone H4) (Herrmann *et al*, 2017; Abdrabou *et al*, 2020; Alanko *et al*, 2015). The results showed that the control proteins were clearly enriched in their corresponding fractions, demonstrating the efficacy of the subcellular fractionation protocol (Fig 1C). In accordance with the mass spectrometry data analysis, we found that TLN1 co-purifies with the nuclear and chromatin-associated fractions of all the cell lines tested.

Next, we evaluated the strength of the TLN1-chromatin interaction by differential salt fractionations. The pellet, which remained after collecting the nuclear fraction, was incubated with increasing concentrations of NaCl (0.3, 0.45, 0.6, 0.8, and 1.2 M) to release proteins bound to the chromatin. Proteins weakly bound to chromatin (e.g. transcription factors) are soluble in low concentrations of NaCl, while proteins tightly bound to chromatin (e.g. histones) are only displaced at high concentrations of NaCl (Herrmann *et al*, 2017). In MCF10A and MDA-MB-231 cells, TLN1 co-eluted with histone H4 at the highest concentrations of the NaCl, whereas in HS578T cells, TLN1 co-eluted with lamin A/C throughout the NaCl gradient (Fig 1D) indicating a weaker interaction with the chromatin. These data indicate that the strength of the TLN1-chromatin interaction varies among cell lines, which may be due to differences in the identity of TLN1-interacting partners in these cells. Taken together, our results clearly demonstrate that TLN1 is localized in the nucleus, where it strongly interacts with the chromatin.

### TLN1 concentrates in specific areas within the nucleus

To investigate the nuclear distribution of TLN1, we performed indirect immunofluorescence staining of TLN1 in HS578T and MDA-MB-231 cells. For this purpose, we used three different antibodies against distinct epitopes of TLN1, one antibody recognizing the head domain and two antibodies recognizing the rod domain, to be able to validate the specificity of the TLN1 fluorescent signal (Fig 2A). The expected localization of TLN1 in the FAs was observed at the bottom plane of the cells, while visualizing the middle plane of the cells displayed nuclear TLN1 signal irrespective of the antibody and the cell line used (Fig 2B-E). Interestingly, a strong nuclear foci pattern was detected in a subset of cells. These foci co-localized with the dark areas of the DAPI staining, which typically correspond to the presence of nucleoli (Di Tomaso *et al*, 2013). Therefore, we co-stained HS578T and MDA-MB-231 cells with an antibody specific for nucleolin (NCL), a well-known nucleolar protein (Jia *et al*, 2017). As anticipated, the immunofluorescent signals corresponding to TLN1 and NCL clearly co-localized, indicating that TLN1 is present in the nucleoli of these cells (Fig 2E). The nucleolar localization of TLN1 is also supported by our mass spectrometry data analysis, which showed 14 nucleolar-associated TLN1-interacting partners with a high confidence score (≥ 0.7 SAINT score) (Fig 1B, left panel) (Table 1).

**Figure 2:**
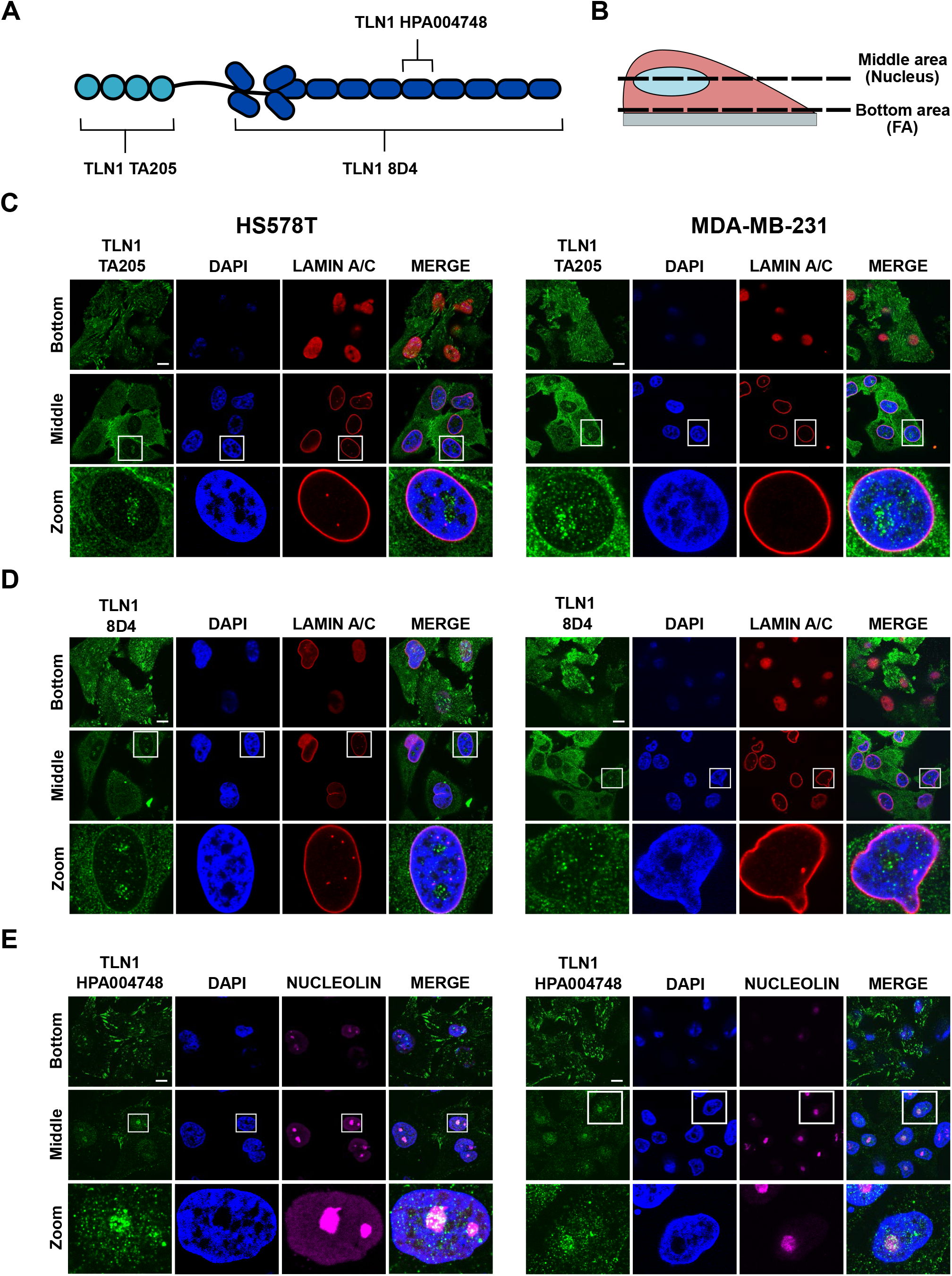
TLN1 concentrates in specific areas within the nucleus. **A)** A schematic overview of the TLN1 epitopes that are recognized by the antibodies used for immunofluorescent stainings. **B)** A schematic overview of single focal planes shown in C-E. Lamin A/C and DAPI were used as nuclear markers, and nucleolin (NCL) as a nucleolar marker. Scale bar 10 µm. Each figure panel shows a different TLN1 immunofluorescent staining performed with the following antibodies **C)** anti-TLN1 TA205 **D)** anti-TLN1 8D4 **E)** anti-TLN1 HPA004748. All images are representative of three biological replicates.

### Nuclear TLN1 regulates gene expression

The nuclear localization of TLN1 and its strong interaction with the chromatin prompted us to investigate whether TLN1 affects gene expression. We depleted TLN1 in HS578T cells by transfecting them with a combination of two small interfering RNAs (siRNAs) specific for TLN1 (siTLN1) and used scramble siRNAs as a control (Scr). The efficacy of the TLN1 downregulation was assessed by immunoblot analysis (Fig 3A-B). Since a prominent reduction in the TLN1 protein levels was detected, we proceeded with analyzing the global gene expression profiles with RNA-seq. Differentially expressed (DE) genes between the Scr and siTLN1 transfected cells were determined with the Bioconductor R package edgeR (Robinson *et al*, 2009), using a log_2_ fold change of at least ± 0.5 and a false discovery rate (FDR) < 0.05 from two biological replicates. Depletion of TLN1 resulted in extensive changes in the gene expression profile of HS578T cells, resulting in the upregulation of 318 genes and the downregulation of 419 genes (Fig 3C) (Table 2). Notably, TLN1 was the most significantly downregulated gene in our RNA-seq screen, thereby confirming the efficacy of the silencing method (Fig 3C, right panel). Normalized gene expression data was used to generate heatmaps from the top 25 upregulated and 25 downregulated DE genes (Fig 3D). Based on the well-known function of TLN1 as a mechanosensitive adaptor protein in FAs, it is plausible that its depletion affected the expression of genes related to cell adhesion. Accordingly, we found several TLN1-dependent genes that are involved in cell adhesion (Fig 3E). Since we found that TLN1 is also present in the nucleus, we searched for genes related to nuclear functions. Among the DE genes, we identified those encoding nuclear structural proteins (NEMP1, H2BC4), chromatin remodelers (BCL7B, MORF4L1), and transcription factors (NFATC2) (Fig 3F).

**Figure 3:**
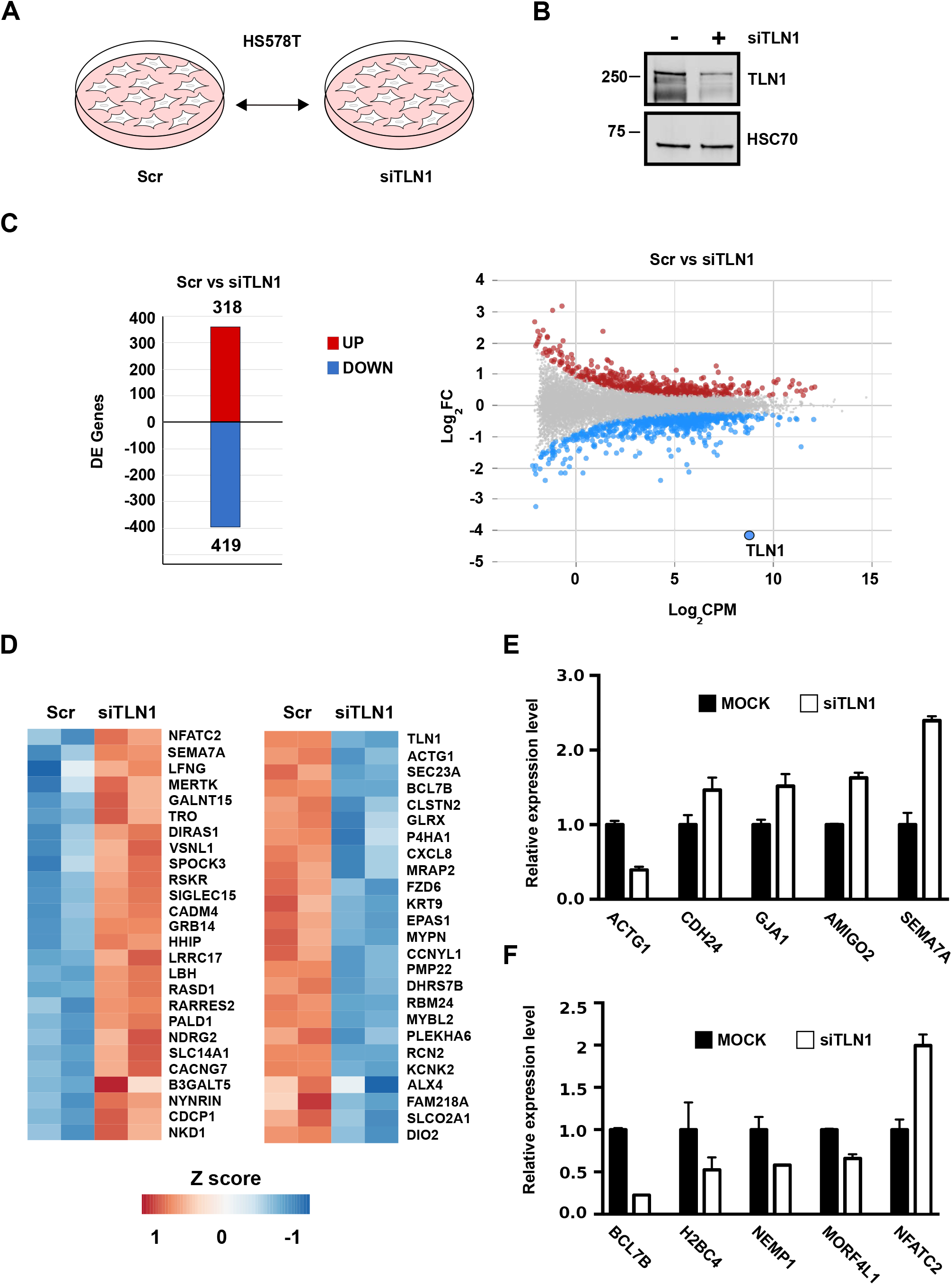
Depletion of TLN1 triggers extensive changes in gene expression. **A)** A schematic overview of the experimental setup for RNA-seq. HS578T cells were transfected with either Scr or siTLN1, and the total RNA from each cell population was extracted and analyzed by RNA-seq. The arrows depict the comparation made for the RNA-seq analysis. **B)** Immunoblot analysis of TLN1 expression in HS578T cells transfected with either Scr or siTLN1. HSP70 was used as a loading control. **C)** Differentially expressed (DE) genes in the Scr vs siTLN1 comparison were determined by Bioconductor R package edgeR (Robinson *et al*, 2009)(Log_2_ FC at least ± 0.5; FDR < 0.05). The number of upregulated and downregulated genes are indicated with red and blue bars, respectively (left panel). Individual DE genes between the Scr and siTLN1 samples were visualized in an MA plot. TLN1 is highlighted (right panel). **D)** The top 50 DE genes were used to generate heatmaps with the CRAN R package pheatmap. The top 25 upregulated and downregulated genes are shown in the left and right panels, respectively. **E-F)** Relative expression levels of five DE genes related to cell adhesion (E) and nuclear functions (F). Error bars +SD. Note that these genes were chosen from the list of total DE genes (Table 2).

These results are, to the best of our knowledge, the first report of a TLN1-dependent gene expression profile. Due to the key role of TLN1 in the FA complexes, it is possible that part of the changes in gene expression upon TLN1 depletion are caused by the disruption of cell-ECM junctions. For instance, it has been shown that the focal adhesion kinase (FAK), a well-established marker of integrin activation, translocates from the cell cortex to the nucleus, where it binds to the chromatin and regulates gene expression (Griffith *et al*, 2021). In addition, the integrity of the FA sites is critical for transforming mechanical stimuli into biochemical signals that eventually affect the expression of different genes (Janota *et al*, 2020). Therefore, it is crucial to distinguish the impact of nuclear TLN1 from the impact of cytoplasmic TLN1 on gene expression.

For interrogating the role of nuclear TLN1 in the regulation of gene expression, we enriched the amount of nuclear TLN1 and determined the mRNA levels of five TLN1-dependent genes: SEMA7A, NFATC2, ACTG1, BCL7B, and SEC23A (Figs 3D and 4B). To specifically increase the level of TLN1 in the nucleus, we constructed a fusion protein consisting of the full-length human TLN1 coupled with GFP and a nuclear localization signal (TLN1-NLS). HS578T cells were transfected with the TLN1-NLS construct, and GFP was used as the corresponding mock control. Analysis with confocal microscopy confirmed that the TLN1-NLS fusion protein was confined inside the nucleus, whereas the GFP control was dispersed in the whole cell (Fig 4A). Next, the mRNA levels of the five TLN1-dependent genes were examined in the mock and TLN1-NLS transfected cells with qRT-PCR (Fig 4B). In line with our RNA-seq data, enriching TLN1 in the nucleus shifted the expression of SEMA7A, NFATC2 and ACTG1 in an opposite pattern to that observed upon TLN1 depletion (Figs 3D and 4B). In contrast, the expression of BCL7B and SEC23A remained unchanged irrespective of the amount of nuclear TLN1, showing that not all TLN1-dependent genes are responsive to nuclear TLN1 (Fig 4B). Taken together, these results demonstrate that nuclear TLN1 regulates the expression of a specific subset of genes.

**Figure 4:**
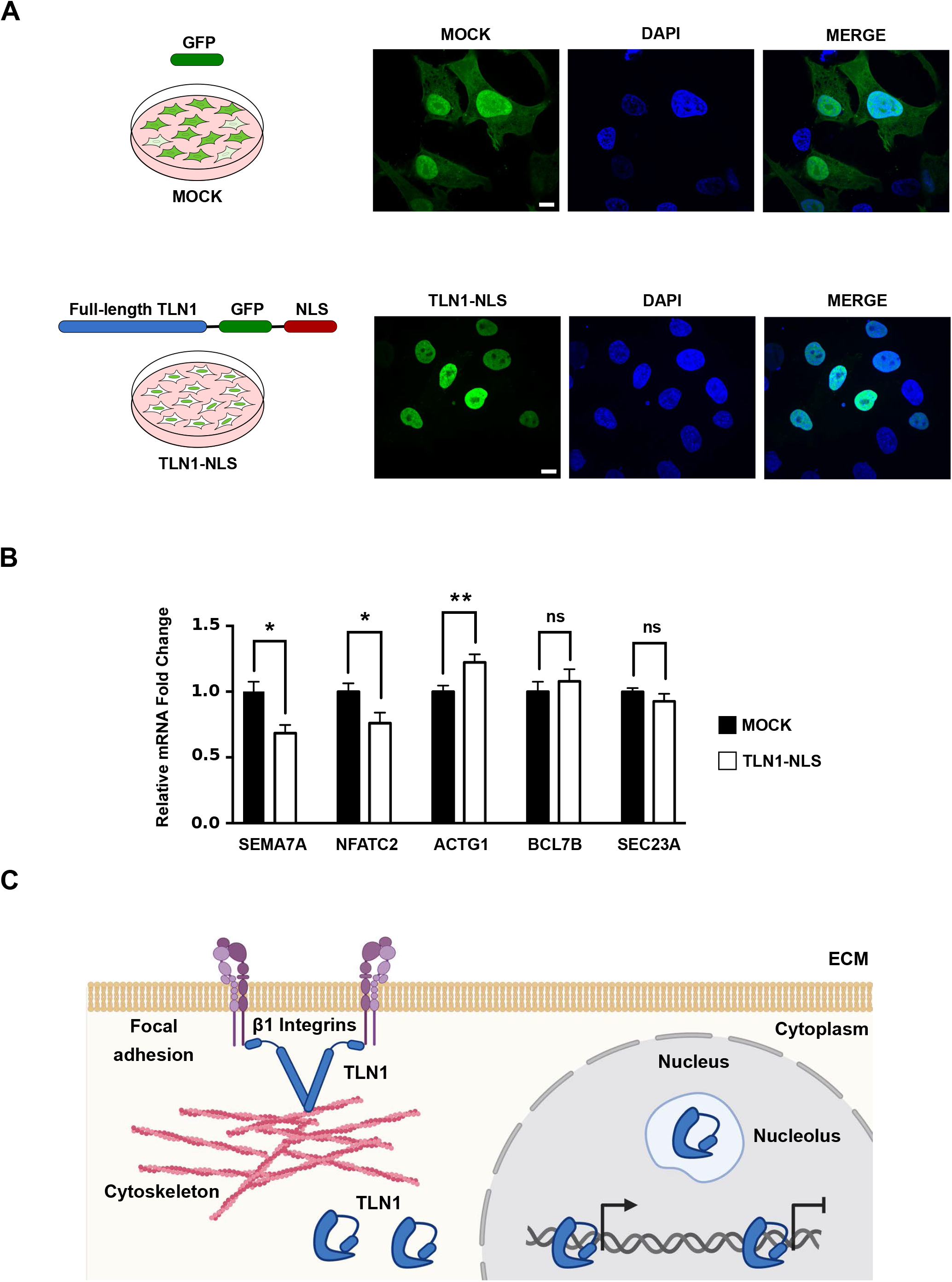
Nuclear TLN1 regulates gene expression. **A)** A schematic overview of the experimental setup to test the role of nuclear TLN1 for gene expression in HS578T cells (left panels). Confocal microscopy images corresponding to maximum intensity projections of the fluorescent signal emitted by GFP or the TLN1-NLS construct (right panels). **B)** mRNA expression of semaphoring 7A (SEMA7A), nuclear factor of activated T cell 2 (NFATC2), actin gamma 1 (ACTG1), BAF chromatin remodeling complex subunit BCL7B (BCL7B), and SEC23 Homolog A (SEC23A) in HS578T cells. The mRNA levels were quantified with qRT-PCR, and GAPDH was used as housekeeping gene. The data is presented as mean values of three biological replicates +SEM, ns: not significant, *p < 0.05, **p < 0.01. **C)** A schematic model of the subcellular localization of TLN1. Apart from being present in the periphery of the plasma membrane and cytoplasm, TLN1 is also present in the nucleus (where it interacts with the chromatin) and nucleolus of the cell. Nuclear TLN1 is represented in close conformation, but this does not exclude the possibility of nuclear TLN1 adopting other conformations. Made with BioRender.com.

## Discussion

The function of TLN1 as a mechanosensitive adaptor protein in integrin adhesion complexes is extensively characterized and continues to be the major focus in the field of TLN1 research. Nevertheless, roles of TLN1 in other cellular compartments have remained enigmatic, which prompted us to investigate the subcellular localization of TLN1. To our surprise, both biochemical and confocal microscopy analyses showed that TLN1 also resides in the nucleus and strongly interacts with the chromatin. RNA-seq analysis revealed that depletion of TLN1 results in extensive changes in the gene expression profile of human breast epithelial cells, which to the best of our knowledge is the first report of gene regulation in a TLN1-dependent manner. Finally, by enriching TLN1 in the nucleus, we demonstrate that TLN1 impacts the expression of a specific subset of the TLN1-dependent genes (Fig 4C).

Active communication between the cell cortex and the nucleus is important to ensure a coordinated transcriptional response upon extracellular stimuli (Zheng & Jiang, 2022). Among the proteins that participate in the FAs, zyxin, paxillin and FAK have been shown to translocate to the nucleus and regulate gene expression (Wang & Gilmore, 2003; Lim *et al*, 2008). Interestingly, studies centered on FAK have led to the hypothesis that FERM domain-containing proteins act as information mediators between the plasma membrane and the nucleus (Frame *et al*, 2010). Our results support this idea and suggest that TLN1 translocate from the cortex to the nucleus to regulate gene expression in response to extracellular stimuli. However, further studies are required to determine the mechanisms mediating TLN1 nuclear translocation.

In the light of our results, it is interesting to address a question how nuclear TLN1 is able to regulate gene expression. Previous studies have shown that nuclear FAK controls chromatin accessibility to allow the binding of transcription factors to specific genomic loci (Griffith *et al*, 2021). In addition, nuclear actin and nuclear myosin-1C, which were initially thought to be strictly cytoplasmic proteins, alter chromatin landscape and also cooperate to maintain active RNA polymerase II at gene promoters (Klages-Mundt *et al*, 2018; Almuzzaini *et al*, 2015). Our results demonstrate that TLN1 strongly interacts with the chromatin of human epithelial cells, suggesting that it could function in a similar manner as nuclear actin, nuclear myosin-1C and/or nuclear FAK. The finding that TLN1 is present in the nucleolus also offers an entry point to exploit the mechanisms by which TLN1 contributes to gene regulation. To this end, it is tempting to speculate that TLN1 plays a role in rRNA maturation, since proteins that are required for the synthesis (e.g. POLR2E, POLR2H, and POLR1C) and maturation (e.g. NIP7 and NOP16) of the polycistronic rRNA precursors were found among the previously identified TLN1-interacting proteins (Tafforeau *et al*, 2013; Goodfellow & Zomerdijk, 2013) (Table 1).

Due to the mechanosensitive role of TLN1 in FA complexes, it is important to consider whether nuclear TLN1 might also respond to mechanical stimuli. Over the last years, it has been recognized that the nucleus reacts to mechanical forces from the extracellular space and the cytoskeleton (Janota *et al*, 2020). Among the best characterized mechanisms of nuclear mechanosensitivity is the protein machinery known as linker of nucleoskeleton and cytoskeleton (LINC) complex (Jahed & Mofrad, 2019). The LINC complex connects the nuclear envelope with the contractile cytoskeleton, and this physical connection mediates the transmission of force to proteins in the inner periphery of the nuclear envelope including nuclear lamins (Miroshnikova *et al*, 2019; Khilan *et al*, 2021). Curiously, the rod domain of TLN1 is composed of 13 α-helical bundles that function as mechanosensitive switches, which change their conformation to expose binding sites for different interacting partners (Goult *et al*, 2021). Thus, one can hypothesize that TLN1 acts as a nuclear mechanosensitive signaling hub. In support of this view, our mass spectrometry data analysis revealed that TLN1 interacts with nine proteins associated with the nuclear envelope, of which emerin mediates changes in nuclear stiffness upon stimulation of the LINC complex (Janota *et al*, 2020). Taken together, we report an unprecedented property of TLN1 by showing that in addition to its association to the plasma membrane and cytoplasm, TLN1 also resides in the nucleus where it interacts with the chromatin and regulates gene expression. These results provide a paradigm shift in the field of TLN1 research, thereby expanding the canonical view of TLN1 subcellular localization and function.

## Materials and methods

### Cell culture

All cells were maintained at 37°C in a humidified 5% CO_2_ atmosphere. MCF10A cells were cultured in DMEM/F12 (Dulbecco’s Modified Eagle’s media, 11330-032, Gibco) medium supplemented with 10 µg/ml cholera toxin, 4% horse serum, 10 μg/ml insulin, 10 μg/ml EGF, 0.5 mg/ml hydrocortisone, and 100 μg/ml penicillin-streptomycin. HS578T cells were cultured in DMEM (Dulbecco’s Modified Eagle’s media, D6171, Sigma-Aldrich) supplemented with 10% fetal calf serum, 2 mM L-glutamine, 100 μg/ml penicillin-streptomycin, and 10 μg/ml insulin. MDA-MB-231 cells were cultured in DMEM (Dulbecco’s Modified Eagle’s media, D6171, Sigma-Aldrich) supplemented with 10% fetal calf serum, 2 mM L-glutamine, and 100 μg/ml penicillin-streptomycin. PC3 cells were cultured in RPMI (Roswell Park Memorial Institute, 1640, Sigma) supplemented with 10% fetal calf serum, 2 mM L-glutamine, and 100 μg/ml penicillin-streptomycin.

### Transfections and gene silencing

All transfections were performed using the NEON Transfection System (MPK5000, Thermo Fisher Scientific) according to the manufacturer’s instructions. Briefly, 2.2×10^6^ HS578T cells were suspended in 100 µl of resuspension buffer, mixed with either 13 µg of DNA or 1.6 µM of RNA, and electroporated using 3×20ms 1050V pulses. To silence TLN1 an equal mixture (1:1) of two siRNAs was used, the siRNAs were purchased from Dharmacon: Cat. No. J-012949-06-0010 and Cat. No. J-012949-07-0010. As a control the ON-TARGETplus Non-targeting control pool from Dharmacon was used (Cat. No. D-001810-10-20). All RNAs were used with a final concentration of 1.6 µM.

### Plasmid construction

The TLN1-NLS plasmid was generated by cloning full-length human TLN1 from PC3 cells. Total RNA was isolated with RNeasy mini kit (74106, QIAGEN) and the complementary DNA was synthetized using random hexamer primers (SO142, Thermo Fisher Scientific). The full-length TLN1 was inserted into a pEGFP-N2 vector using the In-Fusion HD cloning kit (Takara Bio USA). Silent mutations on TLN1 were generated with the QuikChange site-directed mutagenesis kit (Agilent Technologies) to make our construct resistant against the previously described siRNA specific for TLN1 (Cat. No. J-012949-06-0010 and Cat. No. J-012949-07-0010, Dharmacon). Finally, a gene strand containing a Strep-tag II and the NLS of c-myc (Dang & Lee, 1988) was purchased from Eurofins Genomics and inserted in the C-termini of GFP with the In-Fusion HD cloning kit (Takara Bio USA). All primers used for the plasmid construction are listed in Table 3.

### Gene Ontology (GO) term analysis

GO term analyses were performed with the online applications ShinyGO (Ge *et al*, 2020). For analysis performed with ShinyGO the following parameters were used: species: human, p-value cutoff: 0.05, and number of top pathways to show: 30.

### Immunoblotting

Cells were washed and collected in PBS (L0615, BioWest). After collection, the cells were lysed in lysis buffer (10% glycerol, 1 mM EDTA, pH 7.4, 150 mM NaCl, 2 mM MgCl_2_, 1% Triton X-100, 50 mM HEPES pH 7.4, 1 x Protease Inhibitor Cocktail [04693159001, Roche Diagnostics], 50 mM NaF, 0.2 mM Na_3_VO_4_). The protein concentration was determined by BCA assay (23225, Thermo Fischer Scientific). Equal amounts of total protein were resolved on 4–20% Mini-PROTEAN® TGX precast gels (Bio-Rad). The proteins were transferred to a nitrocellulose membrane, which was blocked in 5% milk-PBS with 0.3% Tween 20 for 1 h at room temperature. The primary antibodies were diluted in 0.5% BSA-PBS-0.02% NaN_3_. The following primary antibodies were used: anti-α tubulin (AB 1157911, Developmental Studies Hybridoma Bank), anti-β_1_ integrin (610468, BD Biosciences), anti-histone H4 (05-858, Millipore), anti-HSC70 (ADI-SPA-815, Enzo Life Sciences), anti-lamin A/C (ab26300, Abcam), anti-TLN1 205 (ab78291, Abcam), and anti-TLN1 8D4 (T3287, Sigma Aldrich). The nitrocellulose membranes were incubated with the primary antibodies overnight at 4°C. Secondary HRP-conjugated antibodies were purchased from Promega or GE Healthcare (anti-mouse Cat. No. W4021, Promega; anti-rabbit Cat. No. W4011, Promega; anti-rat Cat. No. NA935V, GE Healthcare). All secondary antibodies were diluted in 5% milk-PBS with 0.3% Tween 20. The nitrocellulose membranes were incubated with the secondary antibodies at least 1 h at room temperature, and then incubated with enhanced chemiluminescence reagent (28980926, GE Healthcare; 34579, Thermo Fisher; 34094; Thermo Fisher). Images were acquired with an iBright imaging system (Thermo Fisher). Unless indicated, all immunoblotting experiments were performed three times.

### Confocal microscopy

For confocal microscopy analyses, 8×10^4^ HS578T or MDA-MB-231 cells were plated on MatTek plates (P35GC-1.5-14-C MatTek corporation) 48 h before imagining. Cells were fixed with 10% paraformaldehyde for 10 min, permeabilized with 0.1% Triton X-100 in PBS and washed three times with PBS. The cells were blocked with 10% FBS-PBS for at least 1 h at room temperature, and then incubated with the corresponding primary antibody dilution overnight at 4^°^C. The following primary antibodies were diluted in 10% FBS-PBS: 1:220 anti-lamin A/C (ab26300, Abcam), 1:100 anti-nucleolin ZN004 (39-6400, Thermo Fischer Scientific), 1:100 anti-TLN1 205 (ab78291, Abcam), 1:100 anti-TLN1 8D4 (T3287, Sigma Aldrich), and 1:100 anti-TLN1 (HPA004748, Sigma Aldrich). After primary antibody incubation the samples were washed three times in PBS and incubated with the corresponding secondary antibody at room temperature for 1 h. The following secondary antibodies were diluted 1:500 in 10% FBS-PBS and used: goat anti-rabbit Alexa Fluor 488 (A11008, Life Technologies), donkey anti-mouse Alexa Fluor 555 (A31570, Life Technologies), goat anti-rabbit Alexa Fluor 546 (A11010, Life Technologies), and goat anti-mouse Alexa Fluor 488 (A11001, Life Technologies). After secondary antibody incubation, the cells were washed in PBS, incubated with 300 nM DAPI diluted in PBS for 5 min, washed again with PBS, and covered with VECTASHIELD (H-1000, Vector Laboratories) mounting medium. Images were captured with a 3i CSU-W1 spinning disc confocal microscope (Intelligent Imaging Innovations).

### Subcellular fractionations

Cells (MCF10A, MDA-MB-231, HS578T, and PC3) were treated with trypsin, collected in culture media, washed with PBS and counted. The 13% of the cell suspension was set apart for preparation of the whole cell lysate using lysis buffer (10% glycerol, 1 mM EDTA, pH 7.4, 150 mM NaCl, 2 mM MgCl_2_, 1% Triton X-100, 50 mM HEPES pH 7.4, 1 × Protease Inhibitor Cocktail [04693159001, Roche Diagnostics], 50 mM NaF, 0.2 mM Na_3_VO_4_). The remaining (87%) cell suspension was used for subcellular fractionation. Cytoplasmic, nuclear, and chromatin fractions were prepared using NE-PER Nuclear and Cytoplasmic Extraction Reagents (78833, Thermo Fisher Scientific) according to manufacturer’s instructions. Briefly, the wet volume of the cell pellet was estimated by considering that 2×10^6^ is equal to 20 µl of wet volume. The cell pellet was suspended in cytoplasmic extraction reagent I, vortexed 15 s, and incubated on ice (see Table 4 for incubation and vortex times per cell line). After incubation, the cytoplasmic extraction reagent II was added. The suspension was incubated on ice and centrifuged (20,000×g, 5 min). The supernatant was collected (cytoplasmic fraction), and the pellet was washed three times with cold PBS. After the washes, the pellet was resuspended in 100 μl of nuclear extraction reagent, incubated on ice and centrifuged (20,000×g, 10 min). The supernatant (nuclear fraction) was collected, and the pellet was washed three times with cold PBS. Following the washes with PBS, the pellet was resuspended in nuclear extraction reagent, supplemented with 1×10^3^ micrococcal nucleases (M02475, New England Biolabs), 5 mM CaCl, and 1 × Protease Inhibitor Cocktail (04693159001, Roche Diagnostics), and incubated at 37°C for 5 min. After incubation the suspension was centrifuged (20,000×g, 10 min) and the supernatant was stored as the chromatin fraction. The protein concentrations of the fractions were determined by BCA assay (23225, Thermo Fischer Scientific).

### Differential salt fractionation

Once the nuclear fraction was collected, the pellet was incubated for 30 min at room temperature with the corresponding dilution of NaCl. NaCl was diluted in water and the following concentrations were used: 0.3, 0.45, 0.6, 0.8, and 1.2 M. After incubation in the first fraction of the NaCl gradient, the suspension was centrifuged (20,000×g, 10 min), the supernatant was collected, and the pellet was incubated in the second fraction. This procedure was repeated consecutively until the pellet was exposed to all the concentrations of NaCl.

### RNA-sequencing

HS578T cells were transfected with either Scr or a combination of two siRNAs specific for TLN1 (Cat. No. J-012949-06-0010 and Cat. No. J-012949-07-0010, Dharmacon). After 48 h the cells were collected, and total RNA was purified with AllPrep DNA/RNA/miRNA Universal Kit (80224, QIAGEN) according to manufacturer’s instructions. The RNA library was prepared according to Illumina stranded RNA preparation guide (1000000124518). Briefly, poly-A containing RNA molecules were purified with poly-T oligo magnetic beads and fragmented with divalent cations under elevated temperatures. For first-strand cDNA synthesis, RNA fragments were copied using reverse transcriptase and random primers. In a second-strand cDNA synthesis, dUTP replaces dTTP to achieve strand specificity. Unique dual indexing adapters were ligated to each sample. The quality and concentration of cDNA samples were analyzed with Advanced Analytical Fragment Analyzer and Bioanalyzer 2100 (Agilent, Santa Clara, CA, USA) and Qubit Fluorometric Quantitation (Life Technologies). Samples were sequenced with NovaSeq 6000 S1 v1.5. All the experimental steps after the RNA extraction were conducted in the Finnish Microarray and Sequencing Center, Turku, Finland. RNA-sequencing was performed from two independent sample series.

FastQC v0.11.9 was used to confirm the quality of the raw reads. The paired-end reads were aligned to the human genome (primary assembly GRCh38, GENCODE) with STAR version 2.7.9a (Dobin *et al*, 2013), using the default settings. The number of read pairs mapped to each genomic feature in release 33 of the GENCODE annotation was determined by featureCounts from subread v2.0.1 (Liao *et al*, 2014). Only read pairs where both ends aligned were counted. Differential gene expression analysis was performed using the Bioconductor R package edgeR v3.34.1. Weakly expressed genes were filtered using filterByExpr defaults, and samples were normalized using the trimmed mean of M-values (TMM) method. The threshold for differentially expressed genes was set to log_2_ fold change of at least ± 0.5 and a false discovery rate (FDR) < 0.05 from two biological replicates. The gene expression data was visualized as an MA-plot, produced by the Bioconductor R package Glimma v2.2.0 (Su *et al*, 2017). Z score transformed log2CPM values were used by the CRAN R package pheatmap v1.0.12 to perform K-Means clustering of the differentially expressed genes and to produce the heat map.

### Quantitative RT-PCR (qRT-PCR)

RNA was isolated using a RNeasy mini kit (74106, QIAGEN) according to the manufacturer’s instructions. For each sample, 1 µg of RNA was reverse transcribed with an iScript cDNA Synthesis Kit (#1708891, Bio-Rad). The SensiFAST SYBR® Hi-ROX kit (Bioline) was used for qRT-PCR that was performed with the QuantStudio 3 Real-Time PCR system (Applied Biosystems, Thermo Fisher Scientific). The mRNA expression levels were normalized against the respective GAPDH expression in each sample. All reactions were run in triplicates from samples derived from three biological replicates. Statistical analyses were performed with GraphPad Prism 7 Software (GraphPad Prism Software, La Jolla California USA, https://www.graphpad.com). The statistical significance was analyzed with paired two-tailed student’s t-test. See Table 5 for primers used in the amplification step.

## Data availability

The original data are available at Gene Expression Omnibus (GEO) database under accession number GSE198191.

## Acknowledgments

We thank all the members of Sistonen laboratory for expert support during the preparation of the manuscript. Imaging was performed at the Cell Imaging Core, Turku Bioscience Centre, University of Turku and Åbo Akademi University. The instruments used in this project belong to the infrastructure of Turku Bioscience Centre. We thank Markku Saari and Jouko Sandholm from the Cell Imaging Core of Turku Bioscience Centre for technical assistance and advice. The Finnish Functional Genomics Centre (Turku Bioscience, University of Turku, Åbo Akademi University and Biocenter Finland) is acknowledged for services, instrumentation, and expertise. This project was funded by The Academy of Finland (L.S.), The Sigrid Jusélius Foundation (L.S.), Cancer Foundation Finland (L.S.), Liv och Hälsa (E.H., L.S.), and Magnus Ehrnrooth Foundation (A.J.D.S.).

## Author contributions

A.J.D.S., E.H. and L.S. designed the research. A.J.D.S., H.S.E.H., M.C.P., J.C.P., and E.H performed experiments. H.S.E.H. analyzed the RNA-seq data. A.J.D.S., H.S.E.H., M.C.P., J.C.P., B.T.G., G.J., E.H., and L.S. interpreted the data. A.J.D.S., E.H. and L.S. wrote the manuscript with the contribution of all the authors.

## Conflict of interest

Authors declare that they do not have competing interests.

## Table Legends

**Table 1:** GO term analyses of TLN1 interacting partners with a SAINT score ≥70 performed with ShinyGO

**Table 2:** Differentially expressed genes between Scr and siTLN1 transfected HS578T cells

**Table 3:** Primers and custom DNA strands used to construct the TLN1-NLS vector

**Table 4:** Duration of incubation (4°C) and vortex steps during subcellular fractionations performed with NE-PER kit (78833, Thermo Fisher Scientific)

**Table 5:** Primers used in qRT-PCR

## Notes

### Competing Interest Statement

The authors have declared no competing interest.

